# Transcriptomic changes across subregions of the primate cerebellum support the evolution of uniquely human behaviors

**DOI:** 10.1101/2025.03.03.641249

**Authors:** Katherine Rickelton, John J. Ely, William D. Hopkins, Patrick R. Hof, Chet C. Sherwood, Amy L. Bauernfeind, Courtney C. Babbitt

## Abstract

**Background:** Compared to other primates, humans display unique behaviors including language and complex tool use. These abilities are made possible in part by the cerebellum. This region of the hindbrain, comprising the flocculus, vermis, and lateral hemispheres, has expanded throughout primate evolution, particularly in great apes. Given the cerebellum’s architecture—differing in connectivity, neuron content, and functions across subregions—examining subregional differences is crucial to understanding its evolutionary trajectory.

**Results:** We performed bulk RNA-seq across samples from six primate species, representing 40-50 million years of evolutionary history, across four subregions of the cerebellum (vermis, flocculus, right lateral hemisphere, left lateral hemisphere). We analyzed changes in gene expression with respect to evolutionary relationships via the Ornstein-Uhlenbeck model and found that, on average, 8.5% of orthologous genes are differentially expressed in humans relative to other non-human primates. Subregion-specific gene expression patterns reveal that the primate lateral hemispheres exhibit significant differences in synaptic activity and glucose metabolism, which in turn are highly implicated in neural processing.

**Conclusions:** This study provides a novel perspective on gene expression divergences across cerebellar subregions in multiple primate species, offering valuable insights into the evolution of this brain structure. Our findings reveal distinct subregional transcriptomic patterns, with the lateral hemispheres emerging as key sites of divergence across the six primate species. The enrichment of genes related to synaptic activity, glucose metabolism, locomotion, and vocalization highlights the cerebellum’s crucial role in supporting the neural complexity underlying uniquely human and other species-specific primate behaviors.

## Background

Compared to other primates, humans exhibit distinct behaviors, such as language production and complex tool use, which involve a network of brain regions including the neocortex, striatum, and the cerebellum (1–4). The cerebellum has become a focus of increasing attention for its role in the evolution of cognition across mammals, particularly in supporting behaviors such as motor coordination, social interaction, and other higher-order cognitive processes (5–7). The cerebellum is involved in the planning and refinement of motor sequences and has been studied for its role in posture, balance, eye movement, and working memory (8–10). Recently, fMRI data in humans has hinted that the cerebellum supports other functions in higher cognition, such as language processing and long-term memory (11, 12).

As a part of the hindbrain, the cerebellum connects with the spinal cord to integrate sensory information from the peripheral nervous system and coordinate motor output within the central nervous system. It undergoes a protracted developmental trajectory, beginning during early stages of embryogenesis and continuing postnatally into adulthood (13). While it represents only about 12% of the mass of the brain, it is estimated that roughly 60-80% of the total neurons in the brain are housed within the cerebellum (14–16). Of those neurons, many subclasses are uniquely located within the cerebellum, including Purkinje neurons and granule cells (17).

The mammalian cerebellum can be anatomically divided into three major subregions: the flocculus, the vermis, and the lateral hemispheres (18). The vermis is responsible for functions in motor control and posture (19, 20). The flocculus, considered to be the oldest structure of the cerebellum phylogenetically, is part of the vestibular system and is involved in balance and coordination (20–22). The lateral hemispheres are implicated in motor control and higher-cognitive processes, such as language in humans (23–25). Importantly, the lateral hemispheres, particularly Crus I and II, are interconnected with the motor and prefrontal regions of the neocortex (26). This level of connectivity is considered a possible explanation as to why the cerebellum may be implicated in aspects of cognition.

The primate lineage shows an overall increase in the size of the cerebellum (27, 28). Although the cerebellum is particularly enlarged in the hominoids (apes), different subregions are variable in their relative size (25). For example, the vermis occupies a much smaller proportion of the cerebellum in humans than in other primates (29). In contrast, the lateral hemispheres are much larger in the lineage leading to humans and chimpanzees, particularly those lobules that are highly connected to the neocortex (30). Relatedly, a coordinated enlargement of the left prefrontal cortex and the right lateral hemisphere in humans has been proposed to be significant for the evolution of tool use (31, 32).

In addition to volumetric studies, previous projects utilizing microarray data, RNA sequencing, and methylation have highlighted the cerebellum as an especially unique region of the brain in terms of its gene expression patterns (33–36). In a dataset surveying 18 different primate species, the cerebellum showed significant differences in expression levels of genes with functions in metabolic processes, neural cell development, and cell signaling (37). Previous single-cell RNA-seq datasets have examined cerebellar cell function in depth and have discovered uniquely human gene expression patterns, highlighting the cerebellum’s role in development and evolution (17, 38).

Studies that have analyzed gene expression differences in the cerebellum amongst primates to date, however, lack regional specificity in their sampling, either failing to report which areas of the structure were sampled or sampling from only a single region (33, 39–42). As the cerebellum is a complex structure composed of multiple subregions of varying connectivity, neuron content, and functions, it is important to characterize differences across subregions to fully understand the evolutionary trajectory of this structure. The level of regional specificity is crucial to detect molecular signals that are spatially limited or less abundant across an entire brain structure. Importantly, as different subregions are associated with different phenotypic traits and behaviors, there may be differences in how these subregions are implicated in development as well as neurodegenerative disease states observed primarily in humans. For instance, it has been suggested that abnormalities in the vermis may be associated with autism spectrum disorder and schizophrenia, while abnormalities of the hemispheres are primarily linked to multiple sclerosis and spinocerebellar ataxia (43–48). To better understand the development of these disease states and why many are rarely observed in non-human primates, it is important to study the cerebellar subregions separately.

Given that the cerebellum houses a majority of the neurons found in the brain, including many granule and Purkinje cells, we hypothesize that there will be differences at the level of synaptic signaling that enhance the local processing of both motor and non-motor tasks. We also hypothesize that primates should show differences in metabolic activity, particular in the processing of glucose, as increase in synaptic densities would require increased glucose uptake and metabolism to support synaptic transmission (49, 50). Together, these differences in synaptic functions and glucose metabolism are expected to give rise to species-specific behaviors.

In this current study, we analyzed differences in gene expression across six primate species, representing 40-50 million years of evolutionary history. We obtained samples from each species in four subregions of the cerebellum to compare regional gene expression in a dataset of primate cerebellar tissue. Our analyses of these datasets reveal gene expression patterns across species that are unique to subregions of the cerebellum, highlighting the importance of subregion sampling to better characterize the evolution, development, and functional roles of the cerebellum.

## Results

### Multidimensional scaling analysis highlights species-specific expression patterns

We analyzed bulk RNA sequences from six primate species (four hominoids, one cercopithecoid, and one platyrrhine with replicates) across four cerebellar subregions (flocculus, vermis, left lateral hemisphere, right lateral hemisphere; Figure 1, Supplementary Table 1). Transcripts were mapped to each pecies reference genome (see Methods), and we quantified the expression of 9,013 total one-to-one orthologs (Supplementary Table 2). To explore the expression differences between these samples, we performed a principal coordinate analysis (PCA) on this dataset. For the total dataset, we plotted the first three axes and found that most of the variation in gene expression can be explained at the level of species phylogenetic relatedness, rather than subregions of the cerebellum (Figure 2A). We found that these PCAs accurately replicate phylogenetic relationships, with more closely related species (*i.e.,* non-human hominoids) clustering together. These phylogenetic relationships are maintained also in PCAs where the dataset is subset by each subregion (Figure 2B). Additionally, we performed an analysis of covariance (ANCOVA) to look at the relative contributions of various factors to gene expression. Species identity and family membership were the most significant sources of variation (Supplementary Figure 3). Importantly, individual identity, age, and sex all share relatively low levels of contribution to variation and so we did not focus on these factors for subsequent analysis.

**Figure 1:**
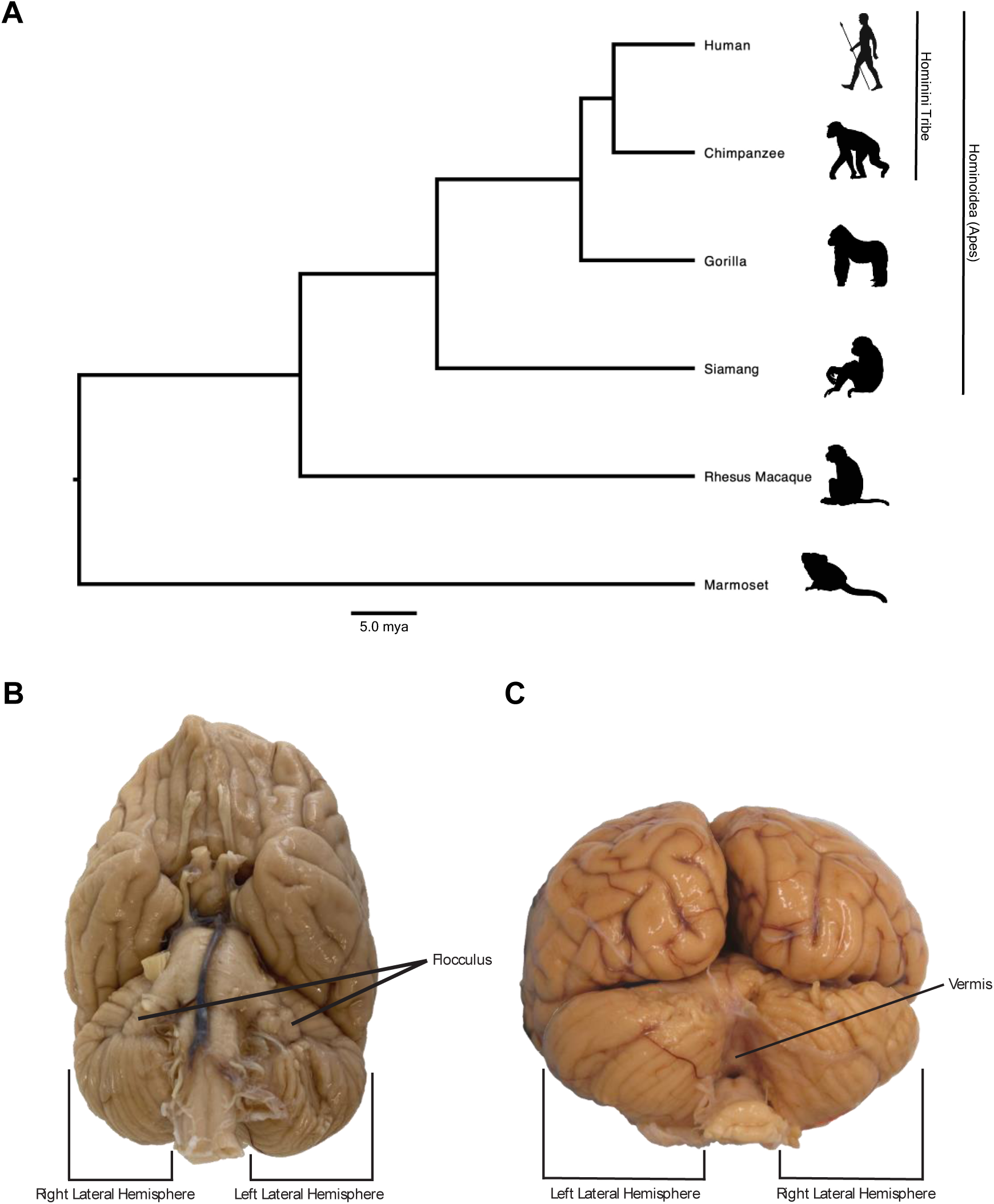
Species and Tissue Sampling Information. A) Primate phylogeny of the five species sampled in this study. The phylogenetic tree is a consensus tree of 1000 iterations produced from 10kTrees v.3 based on GenBank data for each species. The scale bar represents 5 million years of evolution for each branch length. Images of each species were obtained from BioRender.com. B-C) Ventral section (B) and Posterior section (C) of a gorilla brain, depicting the cerebellum. Visible subregions of interest are labeled accordingly. Images courtesy of Chet Sherwood.

**Figure 2:**
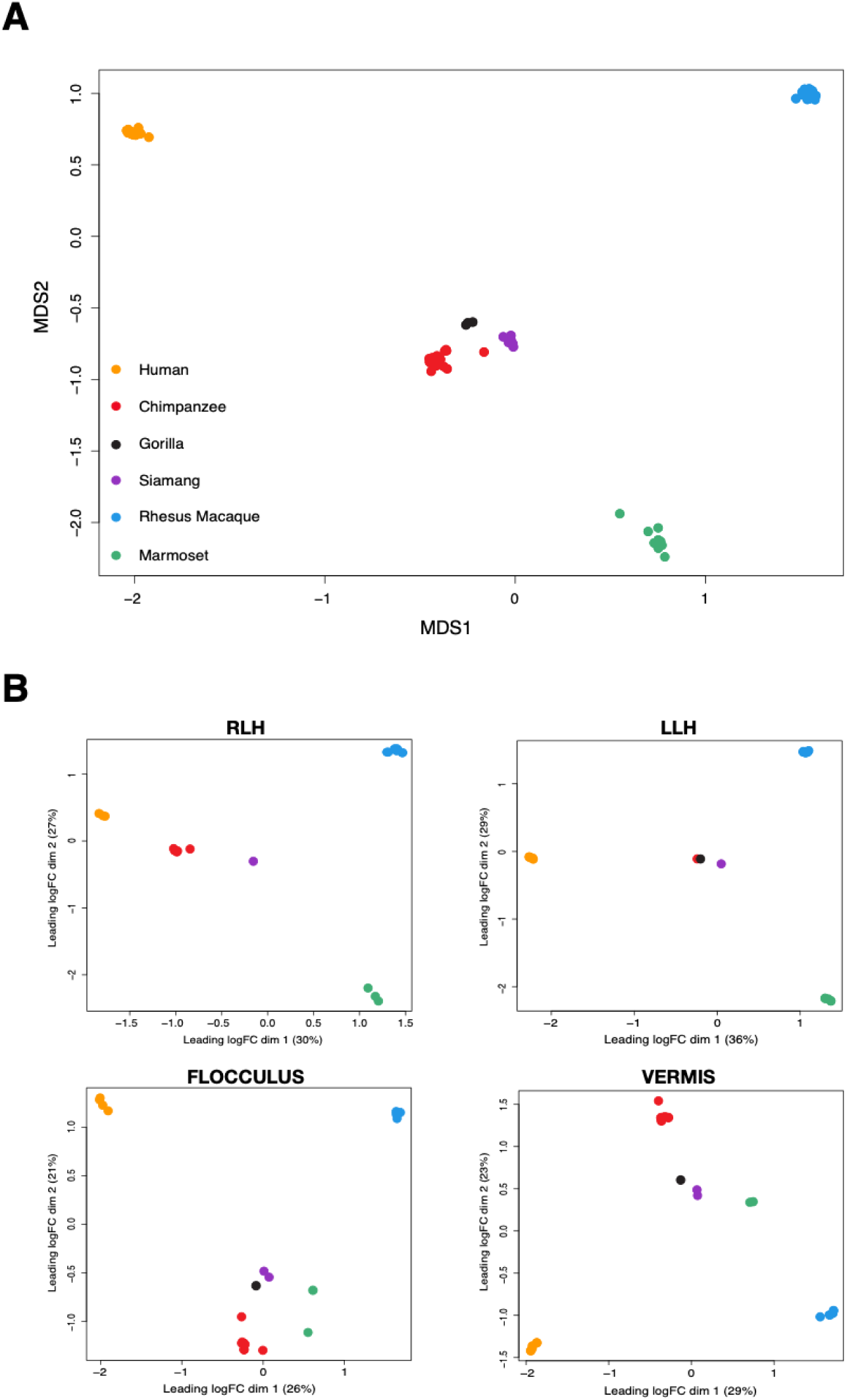
MDS plots of the top 500 DE genes across the primate cerebellum. The first two dimensions are depicted in each plot (percent contribution of each eigenvalue/dimension are shown as axis labels in B). Each plot is based on the top 500 differentially expressed genes as determined by EdgeR. Samples in each plot are color-coded to indicate taxa (key in A). A) All samples. B) Samples are broken down into separate MDS plots by subregion (RLH = Right Lateral Hemisphere, LLH = Left Lateral Hemisphere).

Humans cluster separately from the other hominoid species, highlighting their unique gene expression patterns. This is true for all subregions, suggesting that humans have unique patterns of gene expression across each region of the cerebellum. Based on this finding, we were particularly interested in how the human samples may be driving the differences observed in this dataset. In order to address this, we also analyzed PCAs in which the human samples were removed (Supplementary Figure 4). We saw that the remaining three ape species (two in the right hemisphere) remained tightly clustered and separated from the marmoset and rhesus macaque samples - recapitulating phylogenetic relationships.

### Gene expression phenograms replicate some, but not all, phylogenetic relationships

To further study both between and within species variation, we built expression phenograms at the level of the total cerebellum and also each cerebellar subregion. To do this, we utilized the ape package in R to build a phylogenetic tree with the minimum-distance neighbor joining function (51, 52).

In the total cerebellum, we observed relatively minimal within-species divergence (Supplementary Figure 1A). Human individuals all originate from the same node on the expression phenogram, as do the marmoset, siamang, and gorilla samples. There is some minor within-species variation observed in the chimpanzee and rhesus macaque samples, suggesting that those species show more sample variation. However, bootstrapping values indicate relatively low confidence at those nodes. At a larger scale, we also observed that all ape sample are linked to the same expression phenogram node, reflecting shared evolutionary history. Interestingly, the marmoset group as sister to the hominoid branch and the rhesus macaque samples appear to be the outgroup species. This contradicts the expected phylogeny based on DNA sequence and suggests that the marmosets may show more similar cerebellar gene expression patterns to apes than the rhesus macaque samples do. It is also possible that this may reflect limited sampling in species beyond the apes. The left hemisphere phenogram properly groups the apes as a monophyletic group, with marmosets and rhesus each at a separate node (Supplementary Figure 1B). But again, the marmosets are shown to be sister to the apes rather than the rhesus macaque. For the right hemisphere, the chimpanzee samples are depicted as the outgroup, with the rhesus and marmoset samples shown as sister species (Supplementary Figure 1C). This pattern is also true in the flocculus and vermis phenograms, suggesting that the chimpanzee samples may be particularly divergent from the other species in this dataset (Supplementary Figure 1D-E).

### Rates of gene expression evolution change over time across species and cerebellar subregions

We utilized the Ornstein-Uhlenbeck (OU) model of trait evolution to calculate expression divergences for each gene in our dataset from every species to a reference species (Supplementary Table 3) (53, 54). Using humans as a reference, we plotted the average expression difference across all genes against evolutionary time for each cerebellar subregion (branch lengths; obtained from 10kTrees (55)) (Supplementary Figure 2, Supplementary Table 3). We concluded that pairwise expression differences increasingly diverge over evolutionary time. Importantly, these rates of change do not saturate with evolutionary time, indicating that stabilizing selection is not the dominant process. Rather, saturation of expression differences likely occurs at a node ancestral to these six primate species (54).

### Differential gene expression increases with phylogenetic distance

We conducted differential expression analysis using EdgeR and EVEE-tools in order to assess how the use of different evolutionary models may affect differential expression data. EdgeR utilizes a negative binomial generalized linear model, which assumes a linear relationship between gene counts and sample covariates (56). EVEE-tools is constructed around the Ornstein-Uhlenbeck (OU) model of continuous trait evolution, which accounts for phylogenetic divergence time to estimate the strengths of genetic drift, stabilizing selection and directional selection on a gene expression dataset (54). For both analyses, we computed numbers of differentially expressed genes (DEGs) across pairwise species and clade comparisons. We examined human-specific pairwise species comparisons as well as clade-level comparisons, such as the Hominin tribe and the Ape clade versus non-ape samples (Supplementary Table 4). At the quantitative level, we observed an increase in the number of differentially expressed genes (q-value > 0.05) with greater evolutionary time (Figure 3A-D; Supplementary Figure 5, Supplementary Table 4). That is, when the species, groups, or clades being compared have more phylogenetic distance between them, there is a larger number of differentially expressed genes. This is expected based on the understanding that, over time, genetic variation accumulates due to combined forces of selection as well as genetic drift. At longer timepoints, genetic drift is able to have a stronger effect, resulting in more variation at the level of gene expression. However, this increase in variation eventually levels off at a point beyond the primate lineage (54).

**Figure 3:**
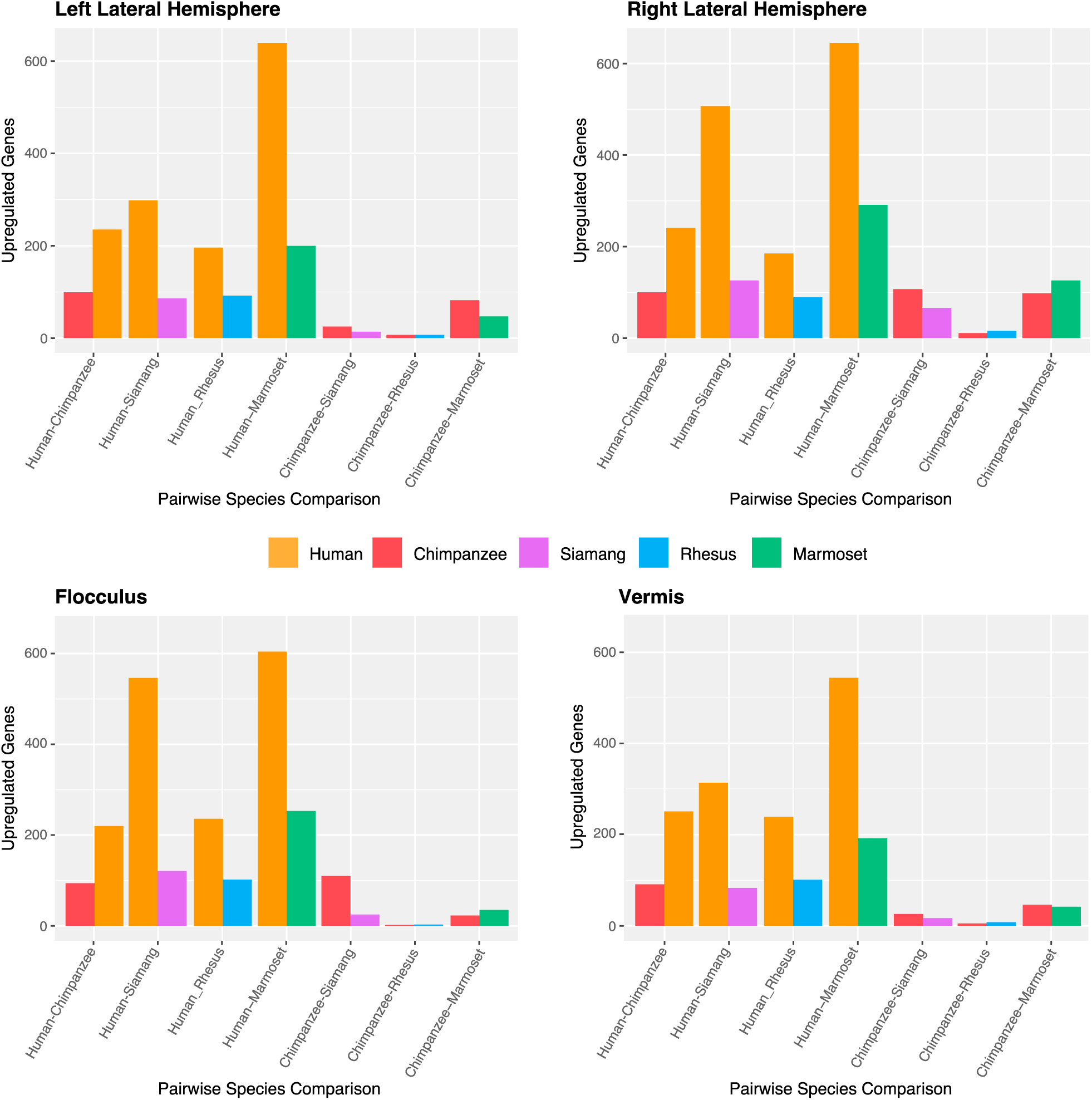
Bar plots of differentially expressed (DE) genes based on pairwise species comparisons. DE genes were determined using the EVEE-tools set of scripts. Only human and chimpanzee pairwise comparisons are shown. Bars are colored according to taxa. Plots are broken down by cerebellar subregion: Left Lateral Hemisphere, Right Lateral Hemisphere, Flocculus, and Vermis.

Upon initial analysis of the EVEE-tools-generated pairwise DEGs, we see that there are not major differences among the cerebellar subregions (Figure 3A-D). The same general trends are shared: comparisons that include human samples show increased levels of DEGs. Unlike what we have observed in other brain regions (37), the comparisons that include chimpanzee samples do not show high levels of differential expression, suggesting that the chimpanzee cerebellum does not appear to be a strong driver of differences in this dataset. Interestingly, human and siamang comparisons show an increase in DEGs that contradicts what we would expect given evolutionary history. We further analyze these DEGs at the qualitative level in sections below, focusing on species and brain-level specific divergences.

#### The lateral hemispheres show divergences enriched in glucose metabolism and synaptic activity

We hypothesized that the left and right lateral hemispheres would show patterns of uniquely human gene expression, with enrichments in categories related to complex motor actions and language processing. Analyzing both the EVEE and EdgeR-generated DEGs, we found significant enrichments in both the left and right lateral hemispheres for categories related to an increase in neural processing (Supplementary Tables 6,7). Genes consistently upregulated in human samples were enriched in categories such as Synaptic Transmission, Dopaminergic Synapses and Presynaptic and Postsynaptic Potential Regulation (Figure 4A; Figure 5D; Supplementary Figure 7, Supplementary Tables 6,7,11). Enrichments in synaptic transmission genes suggest the increased connectivity of the human lateral hemispheres, such that may be supportive of uniquely human behaviors (3, 57). Included in these enriched genes were GABA receptors GABRA4 and GABRA2 as well as the SNCA gene that encodes for α-synuclein, a regulator of neurotransmitter release that is associated with Parkinson’s disease (Figure 5D; Supplementary Figure 7) (58–60). In the human samples, we also observed enrichments for other factors important to synaptic plasticity and neurotransmitter signaling, such as Axon Guidance, Calcium-dependent Signaling, and TGF-β signaling (Supplementary Tables 6,7) (61, 62).

**Figure 4:**
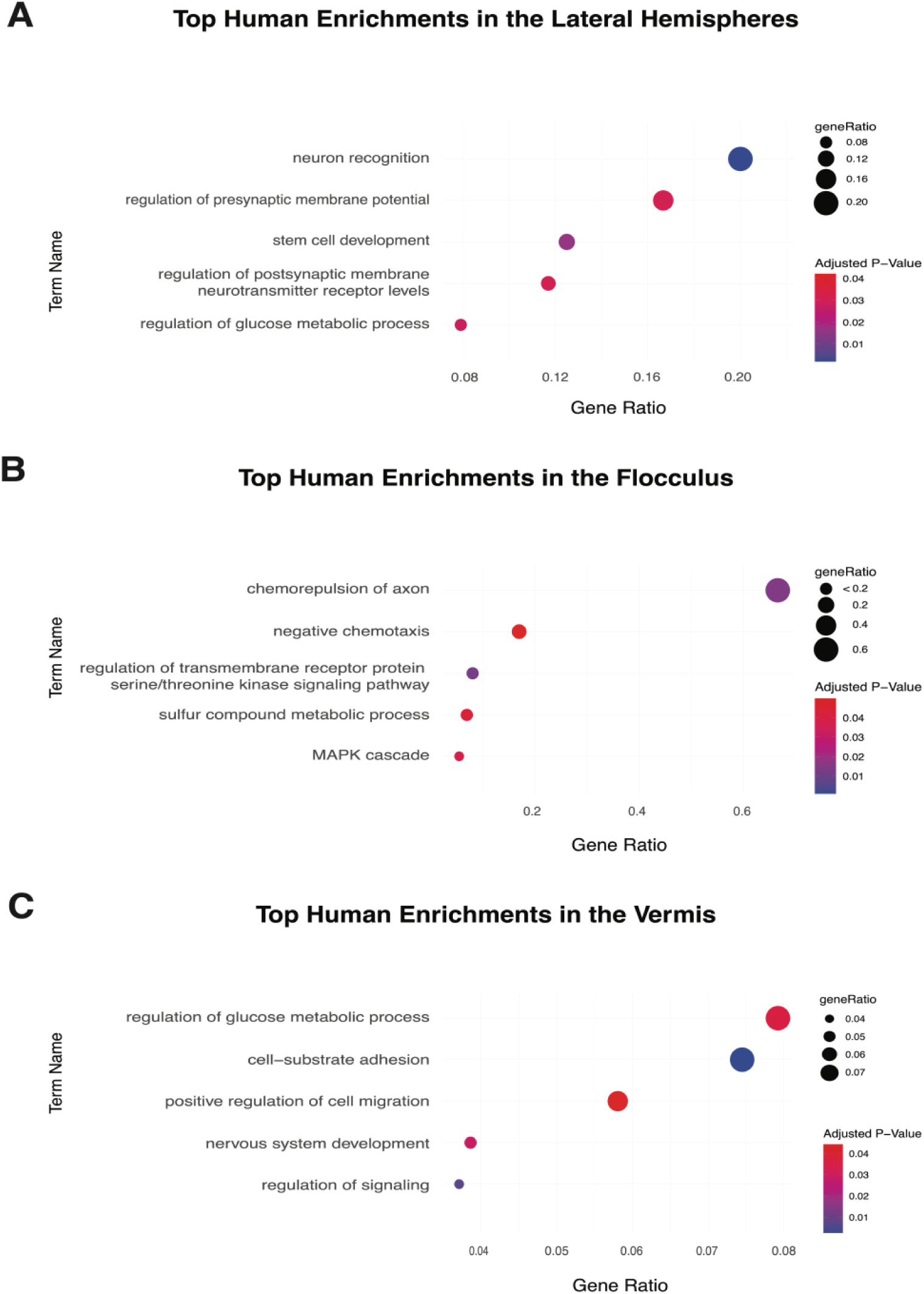
Select human enriched Gene Ontology categories by cerebellar subregion. Terms represented are those significantly (q < 0.05) enriched in human samples compared to non-human primates. Size indicates the ratio of total genes represented by a given GO term, and the color indicates the adjusted P-value (red: lowly significant, blue: highly significant). Placement along the x-axis also represents the ratio of total genes that are members of the GO term. The GO:BP terms were determined by categorical enrichment analysis using GProfiler. A) Top human enrichment categories in the lateral hemispheres. The right and left lateral hemispheres were combined due to significant overlap in enrichment terms. B) Top human enrichment categories in the flocculus. C) Top human enrichment categories in the vermis.

**Figure 5:**
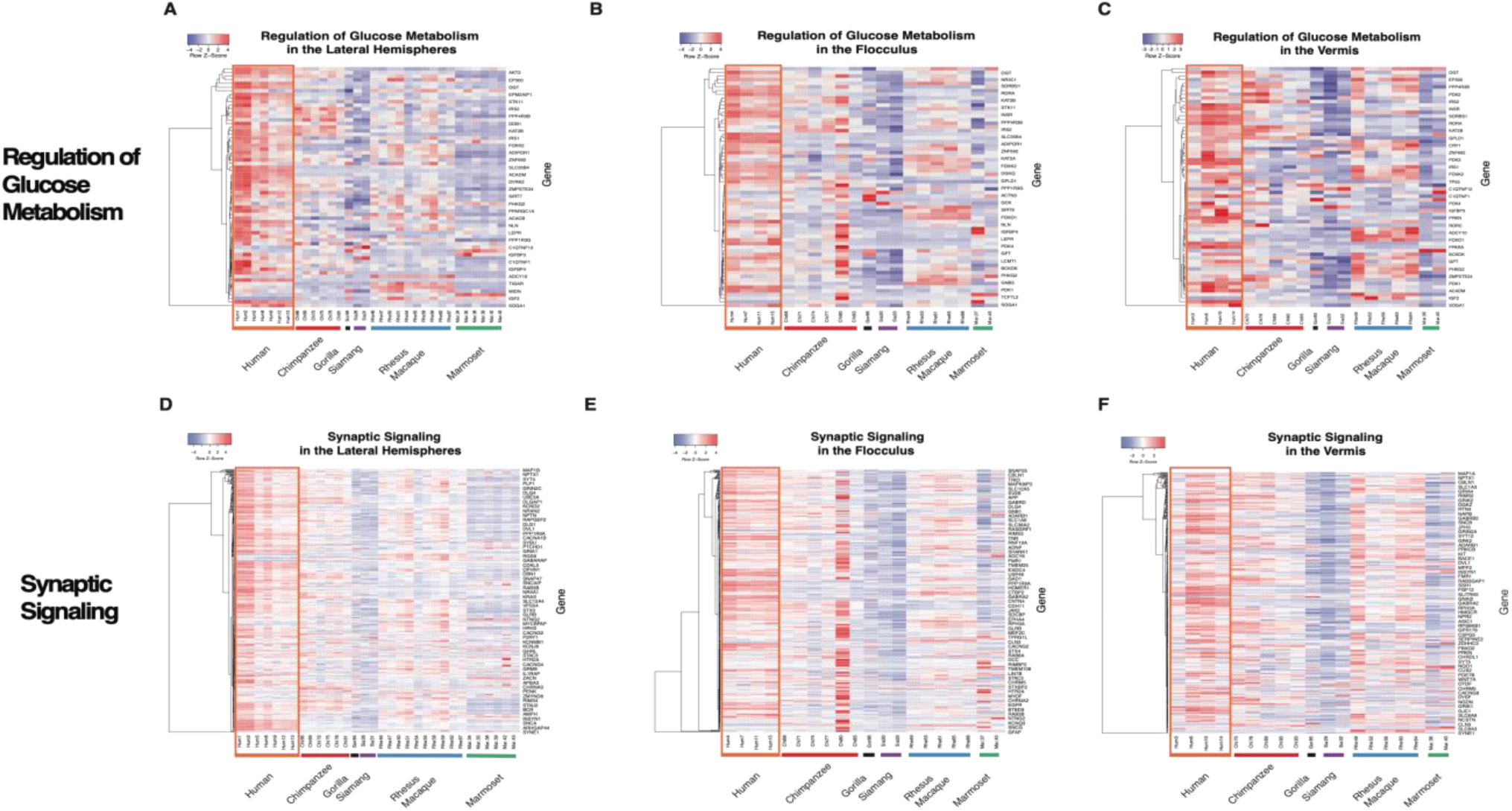
Heatmaps of select human enriched Gene Ontology (GO) categories by cerebellar subregion. A-C) Heatmaps of relative expression for genes that are members of the “Regulation of Glucose Metabolic Process” gene ontology category (GO:0010906). D-F) Heatmaps of relative expression for genes that are members of the “Synaptic Signaling” gene ontology category (GO:0099536). A and D show expression of the left and right lateral hemispheres combined. B and E show gene expression for the flocculus samples, and C and F for the vermis. Each row represents a different gene, and each column is a sample. Samples of the same species are grouped together: labels and color blocks corresponding to species are along the x-axis. The human samples are highlighted by an orange box for emphasis on human-specific expression patterns. Heatmap colors represent relative expression levels (red: more highly expressed; blue: more lowly expressed). All heatmaps were generated using heatmapper.ca (Babicka et al. 2016).

There was a significant enrichment in human lateral cerebellum for glucose-related metabolic processes (Figure 4A; Figure 5A; Supplementary Tables 6,7,10). This is of particular interest as glucose is required to support synaptic activity, and uniquely human upregulation of glucose metabolism is associated with regulation of synaptic transmission (63, 64). One enriched metabolic gene, FOXK1, produces a transcription factor that can induce aerobic glycolysis (65). *PDK3*, another gene enriched in human lateral subregions, encodes an isozyme that catalyzes pyruvate into acetyl-CoA and thus facilitates the transition from glycolysis and the tricarboxylic acid cycle (66). When comparing the two lateral hemispheres, there were higher levels of expression in glucose metabolism-related genes in the right lateral hemisphere observed in the left (Supplementary Figure 6, Supplementary Table 10). This is expected as the right lateral hemisphere has greater connectivity with areas of the left prefrontal cortex in humans, again highlighting the increased neural processing capacity of this tissue (26, 67, 68). However, using GSEA, this difference was not statistically significant and is likely limited by the number of samples available for each region.

Beyond DEGs unique only to humans, we also analyzed differential expression at the clade level to determine what traits may be shared across the ape cerebellum. First, we grouped the human and chimpanzee samples together to represent the Hominin tribe. We observed enrichments in Neural Projection Organization, WNT signaling, MAPK signaling, other non-glucose metabolic pathways, synaptic signaling and axon guidance (Supplementary Table 6). Importantly, we found that synaptic signaling is enriched in this tribe, which suggests that, rather than being unique to only humans, many of these genes were likely upregulated in the ancestor to the human and chimpanzee lineage (Supplementary Table 6) (69). Looking at the apes’ lateral subregions compared with the other two non-ape species, we found enrichments in categories related to Nervous System Development, Neurogenesis, glycogen usage, and non-glucose metabolic processes (Supplementary Table 6).

#### The flocculus and vermis show more conserved metabolic and synaptic signaling functions

In contrast to the lateral hemispheres, the flocculus and vermis have more connections involving the brainstem and spinal cord, and therefore have been primarily characterized for their roles in balance and posture (20, 67, 70). Additionally, the flocculus and vermis show less variability in terms of size across primates compared to the lateral subregions (29). Given these factors, we hypothesized that the flocculus and vermis would show more similar levels of gene expression in genes involved with synaptic activity and glucose metabolism reflecting their conserved function.

Interestingly, in flocculus pairwise species comparisons, humans were enriched for genes related to Synaptic Signaling, Axon Guidance, Neuron Projection Development, and BDNF Signaling (Figure 4B; Figure 5E; Supplementary Tables 6,7, and 11). Brain-derived neurotrophic factor (BDNF) signaling has been studied to encourage synaptogenesis (formation of connections between neurons) and the pathway has been implicated in long-term memory and cognitive processes in humans (71). Also, apes share enrichments in the flocculus for genes involved in Glutamatergic Synapse Development and Maintenance, Central Nervous System Development, Neurogenesis, Cell Projection Regulation, Synapse Organization, and Axonogenesis (Figure 4, Supplementary Table 6), indicating potential roles of the flocculus in higher-cognitive processes supported by enhanced synaptic plasticity and neuron activity.

The vermis was more conserved in terms of gene expression patterning. In the pairwise species comparisons, we observed very few significant enrichment categories (q-value > 0.05) for the DEGs (Supplementary Table 6). Of those that were significant, none appeared to be directly involved in synaptic function (Figure 4C; Figure 5F). For example, the human vermis was enriched for enhanced expression in genes involved in Oxytocin Signaling and Netrin-1 signaling, which highlight the region’s roles in social-emotional brain function and CNS development respectively (72, 73). In contrast to the flocculus, the human vermis was significantly enriched for Regulation of Glucose Metabolism compared to the chimpanzee vermis (Figure 4C; Figure 5C; Supplementary Tables 6 and 7). This is unexpected given the overall highly conserved gene expression patterns observed for the vermis, but points to potential target genes for future studies to understand the functions of this region better. Importantly, this enrichment was not observed in any other pairwise species or clade-level comparisons which suggests that in most primates, glucose metabolism in the vermis is conserved. This study is still limited by the number of subregion tissue samples for each species, and so greater sampling should allow us to elucidate this further.

#### Siamang-specific enrichment patterns across subregions

In the human-siamang comparisons, we observed unexpectedly high levels of differential expression that exceeded that of the more distant rhesus macaque comparisons. As siamangs are only separated from humans by about 15-20 million years (relatively recent compared to the rhesus macaque at 25-30 million years), we would expect fewer DEGs to accumulate due to limited effects from genetic drift (74). However, observing more DEGs suggests that there had been selection pressures that we had not hypothesized.

Investigating this further, we considered the fact that the cerebellum is the primary structure of the brain responsible for regulating motor activity and locomotion (75). For siamangs, this is particularly relevant as they show unique motor behaviors compared to other ape species. Siamangs utilize brachiation, a form of arboreal locomotion in which an animal moves from tree to tree by swinging relying solely on its arms (76). While human and siamang comparisons do not point to any of these genes being specifically involved in motor behaviors, we observed that comparisons between the siamang and the hominin tribe showed significant enrichments in locomotory exploration behavior (GO:0035641) in the right lateral hemisphere (Supplementary Table 6). This gene ontology term describes genes that aid in an organism’s movement as a response to the environment. As the primary habitat of the siamang is the high rainforest canopy, they evolved longer arms to support brachiation as their primary form of locomotion (76). Thus, increased expression in these genes supports this uniquely evolved locomotor behavior.

Compared to the hominin tribe, the siamang vermis and left lateral hemisphere samples were also enriched in genes involved in vocalization behavior (GO:0071625) (Supplementary Table 6). Importantly, siamangs and their close relatives, the gibbons, exhibit unique vocalizations for territorial communication and pair bonding (77). These “songs” are a specialized form of communication is not observed in any other apes (78). One of the mostly highly expressed genes in the siamang cerebellum is *SHANK3*, which has been studied in humans for the role it plays in the development of autism spectrum disorder, particularly how mutations in *SHANK3* are linked to neuromotor deficits and nonverbal behaviors (79, 80).

We therefore hypothesize that the increased number of DEGs found in the siamang comparisons of this study is related to the unique behaviors regarding vocalization and brachiation. This, along with gene expression data that supports cerebellar enrichment in the regulation of synaptic activity, suggests that the cerebellum plays a more important role in language processing and vocal communication than has been previously considered. It also highlights the significant role that the cerebellum has played in primate evolution, particularly in how siamangs have developed.

### Gene co-expression networks highlight regional uniqueness of the primate cerebellum

Further to characterize subregion-specific divergences among the primates in our dataset, we performed a weighted gene co-expression network analysis (WGCNA) (81, 82). This analysis was performed on four of the six primate species—human, chimpanzee, rhesus macaque, and marmoset—to identify groups (or “modules”) of genes with highly-correlated expression patterns. Each module was assigned to a cerebellar subregion based on eigengene values via ANOVA testing (Supplementary Table 8). We compared the number of gene co-expression modules across subregions (Supplementary Figure 9). Having more groups of co-expressed genes indicates that gene expression is highly organized into specific pathways and patterns, suggesting that these regions are more complex and specialized for particular functions. Consistently, we observed a trend across species where the vermis was characterized by relatively few co-expression modules while the flocculus was assigned the most unique modules (Supplementary Figure 9). The hemispheres were much more variable across species in terms of the number of unique expression modules, which is expected based on the differences in pairwise expression data mentioned previously. The flocculus was consistently associated with the highest number of unique modules, likely because of this subregion’s unique connection to the vestibular system and its unique functions in balance and posture (70).

We performed a module preservation analysis to determine whether or not the same modules of genes were conserved across species (83, 84). We focused on the human modules and sought to determine whether the same genes were characterized in similar networks across species. Overall, most identified co-expression network modules are not conserved across species. Of the 30 total co-expression modules observed in humans, only 8 are preserved in chimpanzee, 6 in rhesus macaque, and 2 in marmoset (Supplementary Table 9). While this follows evolutionary divergence patterns, it also suggests that human gene expression modules are largely unique. Of these preserved modules, the left lateral hemisphere showed the highest level of uniqueness with zero conserved modules while the vermis was the most conserved with on average 40% of modules shared amongst species.

We further characterized these preserved networks by performing a hub gene enrichment analysis, where the hub gene is defined as the gene with the most connections to other genes in a co-expression network (82). In the vermis, two hub genes for independent preserved modules are *PPFIBP1* and *CREBBP* (Figure 6B). These are involved in axon guidance and long-term potentiation respectively and suggests that the vermis may have some conserved function in long-term memory processes (85, 86). Additionally, we looked at hub genes for networks that were least preserved and therefore that may signal interesting evolutionary trajectories in the lineage leading to humans. In the left lateral hemisphere, characterized by the least preserved modules, *PLXNA3* is one hub gene (Figure 6A). *PLXNA3* encodes a Semaphorin receptor involved in CNS development by regulating the migration of sympathetic neurons (87). Importantly, *PLXNA3* has been implicated in several neurodevelopmental disorders such as schizophrenia and autism spectrum disorder, suggesting a uniquely human susceptibility to these conditions (88). Further analyses of these hub genes as well as others may aid in our understanding of biological pathway preservation and how this may explain the different evolutionary trajectories of the primate cerebellum.

**Figure 6:**
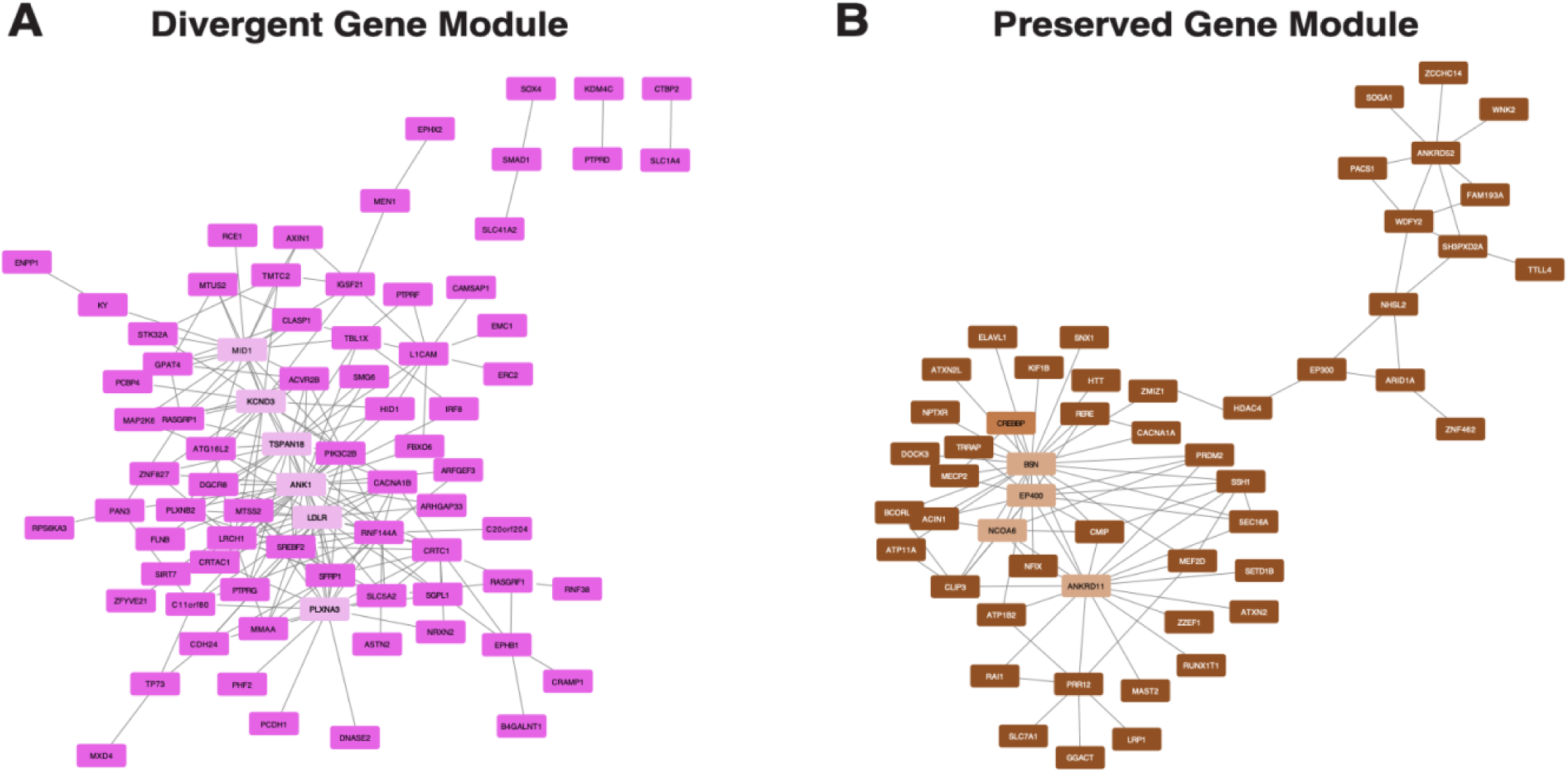
Human modules of co-expressed genes determined by WGCNA analysis. A) Dark Magenta module of genes found in the human left lateral hemisphere depicted as a co-expression network. Module preservation analysis (see Methods) also reveals this module to be highly divergent between humans and other non-human primates for this brain region. Lines connect genes with similar expression patterns. B) Brown module of genes found in the human vermis depicted as a co-expression network. Module preservation analysis finds this module of co-expressed genes to be highly preserved between primate species in the vermis. In both plots, lighter colored-genes are “hub genes”: those with the most connections, therefore driving the co-expression network.

### Extracellular matrix organization distinguishes hominoids from other primates

Across all cerebellar subregions, we observed a consistent upregulation of genes involved in extracellular matrix (ECM) organization in the ape species. The ECM is a large network of connected molecules (mainly proteins) that provides support to surrounding cells (89). In the brain, the ECM has been studied extensively for its involvement in neural cell differentiation, axon guidance, and synaptic plasticity (90). Significantly, the ECM has been characterized as one of the central elements involved in neocortical expansion in mammalian brains (91–93). Studies focusing on the role of the ECM in the cerebellum are limited, but as this region of the brain is also differentially expanded in primates, it is likely that the cerebellar ECM is also a key factor in the cerebellum development among primates (94). Regulation of the ECM is also of interest in studies of neurological diseases, as its role in maintaining tissue homeostasis is often altered and implicated in neuropathology (89). Using these DEGs, future research can study the specific proteins and properties of the cerebellar ECM that differ amongst primates, and specifically how these elements may lead to functional differences.

### Uniquely human enrichments for glioma and glioblastoma-linked genes

In our analyses of uniquely human traits, we were also motivated to analyze disease-relevant genes and how those may differ across the primates sampled in this study. Gliomas and glioblastomas are subsets of brain tumors that are found in the cerebellum (95). Glioblastoma is one of the most aggressive forms of brain cancer, associated with a very poor prognosis in humans (96). Importantly, while these brain tumors have been observed in non-human animals, they are relatively rare in comparison (97). To investigate this difference further, we performed Gene Set Enrichment Analysis (GSEA) to investigate whether glioma and glioblastoma-related genes were enriched in humans and any of the cerebellar subregions included in this study (Supplementary Table 12) (98). When comparing the human samples to all other primate species, we found significant enrichments for glioma-related genes in the right lateral hemisphere (Figure 7). Some of these genes found to be enriched in human samples included DNA mismatch repair proteins *MSH2*, *MSH6*, and *MLH1*. Expression of these genes has been previously investigated for glioblastoma risk assessment (99, 100). While these same gene sets were enriched in the human left lateral hemisphere, flocculus, and vermis, none were statistically significant (q-value < 0.05; Figure 7). This highlights the potential role of the cerebellar right lateral hemispheres in determining susceptibility and prognosis for glioma-related cancers, particularly in humans. This GSEA analysis emphasizes the uniquely human gene expression patterns relevant to gliomas and presents potential evolutionary explanations for the development of these tumors. Importantly, the genes implicated in this analysis may be further studied and potentially serve as treatment targets for clinical research.

**Figure 7:**
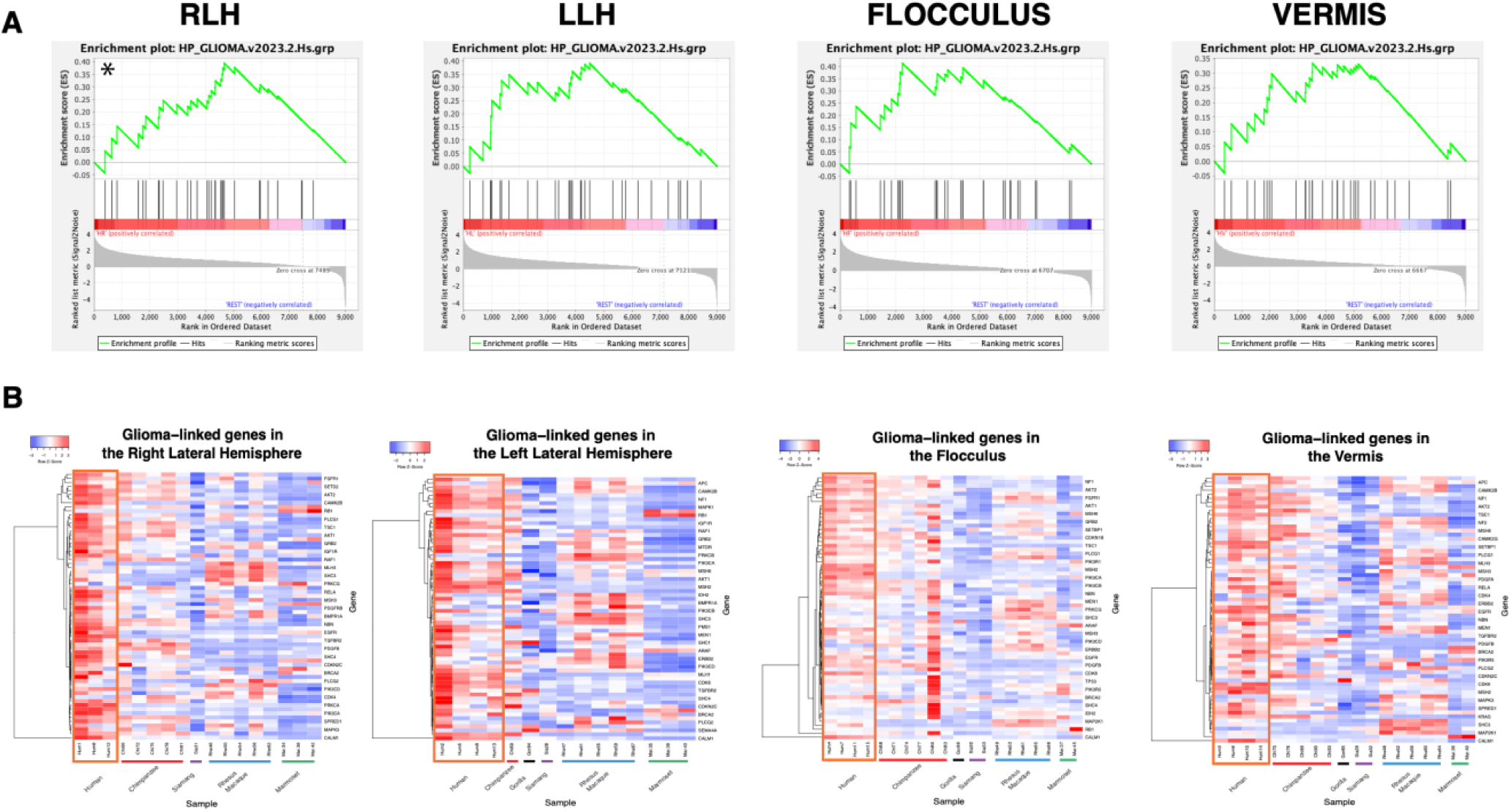
Human-specific enrichments for genes implicated in gliomas across cerebellar subregions. A) GSEA enrichment plots for genes with membership in the HP_Glioma category (HP:0009733). The green line represents the running enrichment score for each gene. The enrichment score is determined by the peak of the curve, and an (*) indicates a significant enrichment score in humans relative to all other non-human primates in this dataset. The middle portion of the GSEA plot ranks the genes based on higher or lower expression in human samples. The bottom portion of the plot shows the value of the ranking metric moving down the list of ranked genes, representing the correlation of gene expression with the phenotype. Each plot represents one of the four cerebellar subregions (RLH = Right Lateral Hemisphere, LLH = Left Lateral Hemisphere). B) Heatmaps of HP:Glioma genes for each of the four cerebellar subregions. Each row represents a different gene, and each column is a sample. Samples of the same species are grouped together: labels and color blocks corresponding to species are along the x-axis. The human samples are highlighted by an orange box for emphasis on human-specific expression patterns. Heatmap colors represent relative expression levels (red: more highly expressed; blue: more lowly expressed). All heatmaps were generated using heatmapper.ca (Babicka et al. 2016).

## Discussion

To characterize the functional role and evolutionary trajectory of the cerebellum better, we analyzed transcriptomic profiles from six primate species, representing roughly 40-50 million years of evolutionary divergence. We compared gene expression differences across the lateral hemispheres, vermis, and flocculus of the cerebellum. We found most of the variation in gene expression can be attributed to phylogenetic relatedness, with other factors, including cerebellar subregion, explaining less variation. Additionally, we found that the number of differentially expressed genes increased over evolutionary time, a reflection of the combined forces of selection and genetic drift. Humans appear to drive a majority of these differences, suggesting the evolution of uniquely human cerebellar functions.

Within each cerebellar subregion, we observed functional enrichments for DEGs that may point to unique specializations. The left and right lateral hemispheres were enriched in processes related to glucose metabolism and synaptic plasticity. As glucose is the primary energy source for the brain, increased glucose metabolic activity suggests that these brain regions may be better able to support higher cognitive processes such as language processing and long-term memory (101). We observed upregulation of glucose metabolism-related genes in humans, suggesting a recent evolutionary shift that corresponds with uniquely human cognitive abilities. Between the two lateral hemispheres, we also observed higher levels of expression in genes related to glucose metabolism in the right lateral cerebellar hemisphere compared to the left. This might be related to the right lateral hemisphere’s connectivity with areas of the left frontal cortex as well as Broca’s area, which is enlarged in humans and highly involved in language production (67, 68, 102). The flocculus was similarly enriched in many synaptic processes and neuronal development genes but did not share the same patterns in glucose metabolic processes as the lateral hemispheres. The vermis, in contrast to all three other subregions, did not show significant enrichments in glucose metabolism or synaptic signaling. The flocculus and vermis are thought to be highly conserved amongst mammals given similar sizes across the primate lineage, lower levels of connectivity to other brain regions, as well as their well-defined roles in vital processes such as motor coordination, balance, and eye movements (20–22, 103). As such, it is not surprising that we did not observe major expression differences across the primate vermis in genes supporting synaptic function and glucose metabolism. However, differences in synaptic processes in the flocculus suggests that this region could possibly be more involved in higher cognitive functions than has been previously assumed.

Much of our analysis focuses on uniquely human expression patterns, but we additionally underscore the magnitude of several non-human primate enrichments. We observed that enrichments in synaptic signaling were shared among humans and chimpanzees, suggesting that some aspects of cerebellar network complexity evolved prior to the splitting of the hominin tribe. The siamang cerebellar samples appeared unexpectedly divergent from other apes, including humans, and many of the differences observed were found to be in genes involved in unique locomotor and vocalization behavior. These differences in gene expression are supported by the unique phenotype of the siamang, characterized by the use of brachiation as well as unique “song” vocalizations for communication. These gene expression divergences help to explain some of the unique characteristics of siamangs, while also highlighting the role that the cerebellum plays in complex behaviors including locomotion and vocal communication. Further sampling of more primate species, including close relatives of the siamang (the gibbons), may help to uncover other interesting roles that this brain region may play in primate evolution.

We also observed enrichments in unexpected processes that warrant further investigation for their roles in cognitive development. We found that extracellular matrix (ECM) organization differed significantly between the hominoids and other primates across all four subregions of the cerebellum. The ECM is a key component of the brain’s microenvironment, providing structural support and influencing cellular behavior (90, 93). The ECM is also known to influence synaptic plasticity, and so differences in the composition of the cerebellar ECM may help explain the behavioral differences across primates. Beyond glucose metabolism, we consistently observed differences in expression across primates for other metabolic processes. In the flocculus and vermis, where glucose metabolism was more conserved, there was an enrichment of DEGs in lipid metabolism, phosphate metabolism, and nitrogen metabolism. While these pathways are not the primary source of energy for the brain, they do point to potentially interesting evolutionary differences within the cerebellum. In particular, lipid metabolism may be of interest for further study as dysfunction in lipid metabolic processes has been linked to cognitive impairment and neurodegeneration, such as in Alzheimer’s disease (104, 105).

We included specific analysis for enrichment of genes related to the development of major brain tumors, such as gliomas and glioblastomas. Genes that have been previously linked to the development of these brain tumors are heavily enriched in human cerebellar tissue compared to other primates, suggesting a potential evolutionary origin for the seemingly increased prevalence of these cancers in humans. Importantly, this analysis does not allow us to assess disease risk using these genes, but rather provides new avenues of research for which genes to target in clinical studies. Looking at how these clinically relevant genes are also differentially expressed in primates may be significant for the success of drug development and the transition between non-human and human clinical trials, as differences across species may affect drug efficacy.

We acknowledge that we only analyze 9013 one-to-one orthologs (roughly one third of total human protein-coding genes) in this dataset. It is therefore likely that observed differences are limited by technological constraints. Additionally, while we have tested and compared two models of estimating differential expression, other potential models may be considered. Using the OU model with only five species is moderately underpowered, and it is possible that other models relying less heavily on long-term evolutionary relationships may be better suited to this dataset. Nevertheless, qualitative enrichments between tested models were similar, suggesting that model choice minimally impacts our conclusions.

Future research should expand our repertoire of species to strengthen evolutionary modeling and statistical power. Other avenues for further research include analyzing these cerebellar subregions at an even deeper level using single-cell RNA sequencing to link the differences in gene expression to cell type identity. Single-nuclei cerebellum datasets have shown that Purkinje cells are responsible for many of the unique gene expression patterns observed in this brain region (17). We analyzed the differential expression of common marker genes for Purkinje cells (marker genes obtained from Sepp, Leiss (17)) and found an overrepresentation in humans compared with the other primate species, suggesting that differences in Purkinje cell expression may be causal to the differences observed across primate species at the bulk level. Further confirming these enrichments with a single cell analysis will ultimately help to determine the significance of this subclass of neurons in primate evolution.

## Conclusion

This study aimed to investigate gene expression patterns in the primate cerebellum regarding the evolution of specialized behaviors in humans, such as language processing, tool use, and complex motor sequences. These behaviors are considered to be cognitively demanding differentiate humans from other closely related great apes. Thus, differences at the level of gene expression in the cerebellum may be informative in understanding the evolution of higher cognition in humans. Our analyses highlight significant gene expression differences between primate species in multiple cerebellar subregions, with many changes in synaptic activity and glucose metabolism, which are highly implicated in neural processing. Direct links between gene expression patterns and higher cognitive capacities remain elusive, however, this study provides mechanisms associated with the evolution of uniquely human traits. This dataset ultimately highlights the uniqueness of the primate cerebellum, suggesting that this brain region may be more involved in higher cognitive functions than has been previously considered.

## Methods

### Biological sample collection and RNA extraction

We obtained cerebellum samples from four human individuals from the National Institute of Health NeuroBioBank. Frozen samples from chimpanzee, gorilla, siamang, rhesus macaque, and marmoset were provided by the National Chimpanzee Brain Resource (www.chimpanzeebrain.org) and various AZA-accredited zoos and primate centers. All individuals were free of any known neurological conditions. Subregion-specific dissections of the vermis, flocculus, and left and right lateral hemispheres (Crus I/II) were performed based on surface morphology. Detailed sampling information is listed in Supplementary Table 1.

All tissue was frozen with a postmortem interval of less than 8 hours. Dissected samples were approximately ∼125 mm^3^ in volume (cubes of tissue approximately 0.5 cm in length on each side) and remained frozen during dissection. Each sample was homogenized using a Tissuelyzer (Qiagen, Hilden, Germany), and total RNA was isolated using an RNAeasy kit (Qiagen) with DnaseI treatment. At least 1 µg of RNA was isolated from each sample for sequencing. The A260/280 (ratio of RNA to protein) was ≥1.7 and A260/230 (ratio of RNA to salts, carbohydrates, and phenol) was ≥2.0. RNA Integrity numbers (RINs) were generated on the Agilent 2100 Bioanalyzer (Agilent Technologies, Santa Clara, California) to assess the quality of the RNA before proceeding to bulk sequencing library preparation.

### Library preparation and RNA sequencing

We constructed single end RNA-Seq libraries using the NEBNext mRNA Library Prep Kit for Illumina (New England Biolabs, Ipswich, MA). Libraries of assorted species and cerebellum subregions were prepared to reduce batch effects. Library sizes and quality were checked on the Agilent 2100 Bioanalyzer (Agilent Technologies). The samples were sequenced on an Illumina HiSeq 3000 (Illumina, San Diego, California) at the Genome Technology Access Center (GTAC) at the Washington University School of Medicine, at a depth of 30 million 50-bp reads.

### Transcriptome assembly and mapping

For mapping, we utilized the following reference genome assemblies available via ENSEMBL: Human (GRCh38), Chimpanzee (Pan_tro_3.0), Gorilla (gorGor4), Gibbon (Nleu_3.0), Macaque (Mmul_10), Bolivian Squirrel Monkey (SaiBol1.0), and Marmoset (Callithrix_jacchus.ASM275486v1). As Siamang and Spider Monkey both do not have annotated references, we intended to map to the next closest available reference for each: Gibbon and Bolivian Squirrel Monkey, respectively. However, the Squirrel Monkey is more closely related to the Marmoset samples than the Spider Monkey and so the Spider Monkey samples were removed for subsequent analysis.

Reads were mapped to orthologs via Bowtie2 (106, 107). HT-Seq was used to count the number of reads per individual sample that map to annotated features in the genome (108). To merge all counts into a single table, the genes for all non-human primate (NHP) individuals were converted into the human orthologs using Blast (NCBI), and only those with 1:1 orthologs across all species were included in downstream analyses. We filtered the dataset to remove genes that showed counts of 0 in more than 20% of the samples in order to remove any mapping inconsistencies found in a single species (Supplementary Table 2). This left us with a total of 9013 one-to-one orthologs for analysis. Furthermore, counts were TMM-normalized in EdgeR to account for variation in library size, length, and GC content (56).

### Distance-based analyses (PCA and phenograms)

We conducted several principal coordinates analyses (PCA) based on pairwise distance matrices of the total dataset as well as each individual cerebellar subregion. We calculated pairwise distances by leading log2 fold change, using previously described methods (38) of the top 500 most variably expressed coding genes. We plotted the PCAs using the plotMDS.DGElist function in the edgeR package in R (56).

We used the same pairwise distance matrix to build phenograms visualizing how similar the samples in the dataset are based on the top 500 most variable genes. We built these trees using the minimum distance neighbor-joining function in the ape package in R and conducted a bootstrap analysis over 1000 iterations (boot.phylo function). This is based on methods originally proposed in Saitou and Nei (52). We created phenograms of the entire dataset and of the four subregions separately.

### Differential expression analysis

As mentioned above, we prefiltered and prenormalized counts prior to differential expression analysis. These analyses were all conducted at the level of each individual cerebellar subregion. To compute differential expression in EdgeR, we performed quasi-likelihood F-tests using the GLM functionality in EdgeR (109). This model assumes that our dataset is nonnormally distributed and is a common model for comparative RNA-seq analysis. Additionally, the quasi-likelihood F-tests are suited to datasets with smaller numbers of replicates. With this, we performed pairwise species and clade comparisons. We compared all of the species in the dataset to the human and chimpanzee samples for a hominin-tribe specific analysis (Species vs Human and Species vs Chimpanzee). We also looked at each species compared to the combined gene expression patterns of the human and chimpanzee samples grouped together (Species vs. Hominins). For clade comparisons, we grouped all of the ape species together (Human, Chimpanzee, Gorilla, and Siamang) and compared these to the non-ape species (Rhesus and Marmoset). With each comparison, we obtained a logFC, logCPM, p-value, and FDR-corrected Q-value for each of the 9013 genes in the dataset. To define a gene as being differentially expressed, we utilized an FDR-corrected Q-value cutoff of 0.05 and a logFC threshold of greater than 1.5 or less than - 1.5.

To compute differential expression using the OU model, we performed analysis utilizing code from the EVEE-tools set of scripts (54). Trees for this analysis were downloaded from 10K trees. We first calculated expression divergence across species in each cerebellar subregion, using the residuals.R script. For these analyses, we tested using different species as the reference species for normalization and found negligible differences in the resulting data. Based on this, we utilized humans as the reference species for all further contrasts. These data are reported and plotted against evolutionary time (extracted from branch lengths of trees) to show the pattern of expression divergence across the cerebellum. We calculated evolutionary means and variance values for each gene across the entire phylogeny, separated by cerebellar subregion, using the fitOUModel.R script, where the q-value shows the significance of fit for the OU model as compared to Brownian Motion (the null model). Differential expression was determined using a multivariate OU model as outlined in the ouRegimes.R script. Regimes were defined by species or clades of interest, similar to the pairwise comparisons computed in EdgeR. This allows us to analyze the differences between the human and chimpanzee samples while also accounting for the shared phylogenetic history of the dataset. P-values were calculated, representing the fit of each OU model against a null Brownian motion model. These P-values were corrected for multiple hypothesis testing using the Benjamini-Hochberg FDR procedure. Additionally, Akaike and Bayesian Information Criterion (AIC and BIC) scores were calculated for each gene in each model. We defined differential expression as those genes with significant AIC and BIC scores against the null, as well as genes that had an FDR-corrected Q-value of < 0.05. To make our differential expression analysis comparable to that calculated in EdgeR, we also computed logFC scores from the normalized counts data and assigned a threshold value of greater than 1.5 and less than -1.5 for genes to be considered differentially expressed.

We conducted differential expression analysis using both of these models and found that overall, fewer DEGs are observed when using the OU model in EVEE-tools in comparison to the GLM model in EdgeR (Supplementary Table 5). This is likely due to the fact that the GLM model employed in EdgeR is a less stringent model and assumes a level of independence in our samples. The EVEE model instead is able to include how the samples in our dataset are phylogenetically related, and as a result it is better able to interpret which genes were more likely to have evolved due to forces of genetic drift or forces of selection. Overall, using both EVEE-tools and EdgeR, we observed similar species level trends as well as subregion-specific trends in the enrichment categories for the lists of DEGs, suggesting that the patterns of expression divergence are conserved regardless of which model is used. For simplicity, we limited our reported differential and enrichment analyses to be based on the EVEE-tools modeling strategy.

### Weighted Gene Correlation Network Analysis (WGCNA)

We used the R package WGCNA in order to generate networks of co-expressed genes in human, chimpanzee, rhesus macaque, and marmoset samples (81, 82). Methods used to generate networks are based on similar analyses (84, 85). With this, we analyzed each species separately with cerebellar subregions combined for normalization to limit batch effects. We normalized RNA-seq data using EdgeR, following similar methods as in our differential expression analysis (56). Weighted networks were calculated using the pickSoftThreshold function. We selected a soft-thresholding power of 14 for consistency across all four species (at this power, all species have an r^2^ value of ≥ 0.8). Following this, we transformed the correlation matrix of gene expression data into an adjacency matrix for each species.

Modules of co-expressed genes were identified through hierarchical clustering and dynamic tree cutting using average linkage values (deepSplit = 2 as a moderately sensitive parameter of splitting up modules). The minimum module size was set to 30 genes for all species. For each module generated, we also calculated the associated eigengene values based on the expression data (Supplementary Table 8).

In order to classify each module to a cerebellar subregion, we performed an ANOVA test to compare expression values across samples (each assigned a subregion) to our eigengenes for each module. We used a p-value cutoff of 0.4 in order to define a module classification as significant, based on natural cutoffs as observed in histograms of the generated p-values (Supplementary Figure 8). We note that, with FDR correction, p-values were largely insignificant, and this is likely a reflection of limited sample sizes for each of the cerebellar subregions. Following module annotation, we obtained a list of the most highly connected genes for each module, known as “hub genes,” using the chooseTopHubInEachModule function in the WGCNA package. These hub genes were analyzed as indicators of overall module function.

We measured the preservation of the human modules across the chimpanzee, rhesus macaque, and marmoset samples in order to better understand the organization of lineage-specific expression changes. We utilized the modulePreservation function in WGCNA with the human module list serving as the reference network. For the other species, we did not include the module assignments so as to ensure consistency and reproducibility across the different species. 8700 genes were compared in this analysis after filtering for those shared amongst all four species, as well as following WGCNA filtering and normalization. We performed 200 permutations of module preservation for each species. A human module was considered preserved in the other species if the Zsummary.pres score was greater than 5 and if the p-value associated with subregion annotation was ≤ 0.4 (similar cutoffs as in [84]). Preserved modules were also analyzed based on hub gene functional enrichments.

### Functional enrichment analysis

To further analyze the functional significance of these differentially expressed genes, we performed several functional enrichment analyses. We utilized gProfiler’s g:OST functional enrichment tool to broadly survey the significantly enriched biological processes (110). This data is reported in Supplementary Tables 6 and 7 (only terms with a q-value > 0.05 are reported). Importantly, we conducted enrichment analyses separately depending on the directionality of the expression in our pairwise comparisons, and thus, were able to characterize which specific processes were upregulated in which specific species or clades of interest. In addition to gProfiler, we also used GSEA to specifically analyze enrichments in glioma and glioblastoma-associated gene sets (98, 111). For this, we downloaded gene sets from the Molecular Signatures Database (MSigDB) (112): KEGG_Glioma.v2023.2, HP_Glioma.v2023.2, and HP_Glioblastoma_Multiforme.v2023.2 (Supplementary Tables 10-12). With these, we again conducted pairwise comparisons at the level of different cerebellar subregions, comparing species to species, clade to clade, and also the human samples to the rest of the species in the dataset. We conducted GSEA analysis with 1000 gene set permutations, and gene sets were considered significantly enriched with an FDR < 25% and a nominal P-value < 0.05 (98). Heatmaps were generated using the GSEA software in order to analyze the expression differences at a gene-specific level.

## Declarations

### Ethics approval

Human samples were exempt from IRB protocols as they are all de-identified. Other non-human primate samples are from the National Chimpanzee Brain Resource.

### Consent for publication

Not applicable.

### Availability of Data

Sequencing data have been deposited in the Short Read Archive: Human reads (https://dataview.ncbi.nlm.nih.gov/object/PRJNA1227155?reviewer=b8cqoa35ei5klaonf82ismhqgv) and Non-Human reads (https://dataview.ncbi.nlm.nih.gov/object/PRJNA1227154?reviewer=sg6624igvsr55el5h8ffr116or).

### Competing Interests

The authors declare that they have no competing interests.

### Funding

National Science Foundation (BCS-1750377; C.C.B.), National Institutes of Health (T32 GM135096; KR), James S. McDonnell Foundation (C.C.S.), National Institutes of Health (NS-092988; C.C.S.), National Science Foundation (SMA-1542848; C.C.S.), and The Leakey Foundation (A.L.B.).

### Author Contributions

*Conceptualization* (C.C.B., A.L.B., C.C.S.)*; Resources* (J.J.E., W.D.H., P.R.H., C.C.S.)*; Methodology* (K.R.)*; Formal analysis* (K.R., C.C.B.)*; Investigation* (C.C.S., A.L.B., C.C.B.)*; Writing – original draft* (K.R., C.C.B.)*; Writing – review and editing* (K.R., J.J.E., W.D.H., P.R.H., C.C.S., A.L.B., C.C.B.)*; Visualization* (K.R.).

## Acknowledgements

We would like to especially acknowledge Mary Ann Raghanti for help with sample allocation. We also thank all members of the Babbitt laboratory for all of their support and feedback. We also thank Drs. Jason Kamilar, ChangHui Pak, Jennifer Rauch, and Shelly Peyton for valuable feedback. Additional thanks to the Dallas Zoo, Busch Gardens Zoo, Tulsa Zoo, and Cleveland Metroparks Zoo for help with sampling (via the National Chimpanzee Brain Resource).

## Notes

### Competing Interest Statement

The authors have declared no competing interest.

