## Supplemental Info and Figures for "Transcriptomic changes across subregions of the primate cerebellum support the evolution of uniquely human behaviors"

**Supplementary Table Information**

**Supplementary Table 1:** Sample Information

**Supplementary Table 2:** Filtered Counts Files (Total, RLH, LLH, Flocculus, and Vermis)

**Supplementary Table 3:** Residuals from EVEE analysis

**Supplementary Table 4:** Differentially Expressed Genes from EVEE analysis

**Supplementary Table 5:** EdgeR vs. EVEE Differentially Expressed genes

**Supplementary Table 6:** Enrichment Data for EVEE DEGs

**Supplementary Table 7:** Enrichment Data for Human Upregulated Genes in EVEE analysis

**Supplementary Table 8:** WGCNA eigengenes

**Supplementary Table 9:** WGCNA Preserved Human Modules

**Supplementary Table 10:** Glucose Metabolism gene expression

**Supplementary Table 11:** Synaptic Signaling gene expression

**Supplementary Table 12:** Glioma-linked gene expression

**Supplementary Figure Information**

**Supplementary Figure 1:** Gene expression phenograms of all samples.

**Supplementary Figure 2:** Mean-square expression differences over evolutionary time for each of the cerebellar regions.

**Supplementary Figure 3:** Boxplots of the distributions of variance contributions as determined through ANCOVA analysis.

**Supplementary Figure 4:** MDS plots by cerebellar subregion of non-human primate samples.

**Supplementary Figure 5:** UpSet Plots of each subregion showing the shared DEGs for all species across the phylogeny.

**Supplementary Figure 6:** Heatmaps of individual lateral hemispheres of genes involved in the regulation of glucose metabolism (GO:0010906).

**Supplementary Figure 7:** Heatmaps of individual lateral hemispheres of genes involved in synaptic signaling (GO:0099536).

**Supplementary Figure 8:** Histogram of p-values from ANOVA test for gene module association with cerebellar subregions.

**Supplementary Figure 9:** Boxplots of different human gene co-expression modules and their relative associations with each cerebellar subregion.

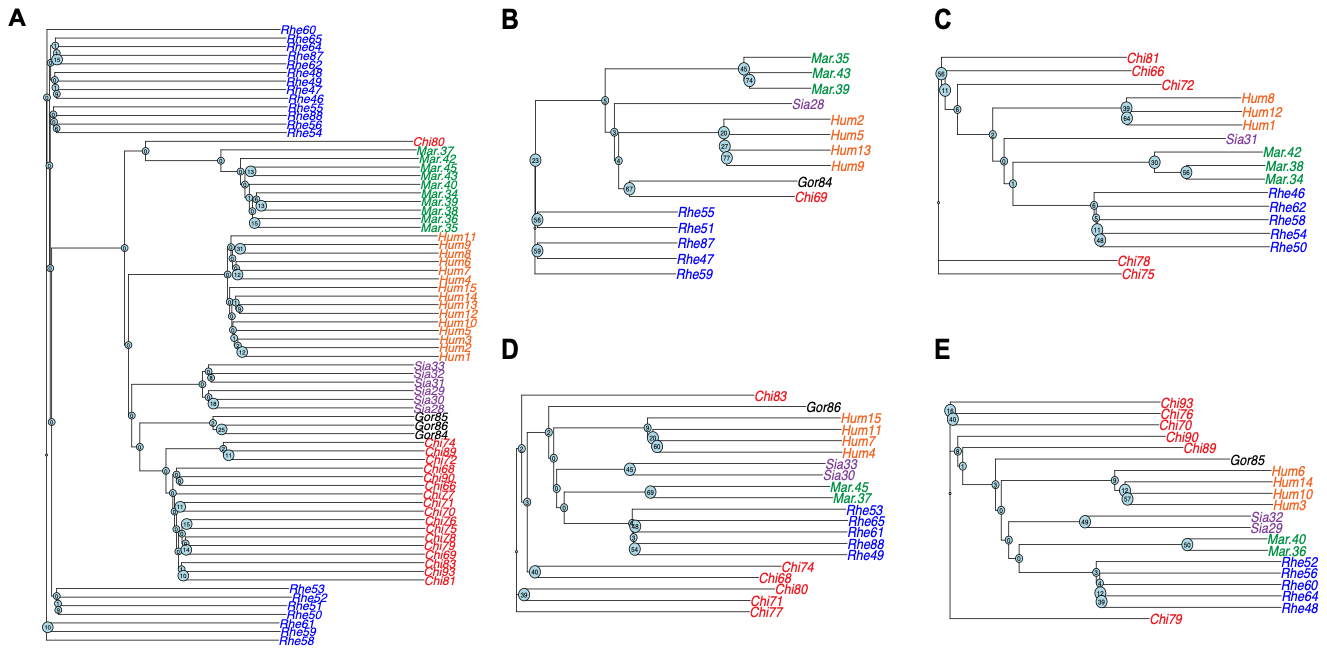

**Supplementary Figure 1: Gene expression phenograms of all samples.** The phenogram was constructed from neighboring-joining tree estimation based on log2 fold-change distances of the 500 most variable genes based on standard deviation. Reproducibility of the nodes of the tree were estimated using a bootstrap analysis of 1000 iterations, with the bootstrap value illustrated at each node. Sample names are color-coded to indicate taxa identity (Blue = rhesus macaque, orange = human, green = common marmoset, purple = siamang, red = chimpanzee, and black = gorilla). A shows the entire dataset, B = Left Lateral Hemisphere (LLH), C = Right Lateral Hemisphere (RLH), D = Flocculus (FL), and E = Vermis (VR).

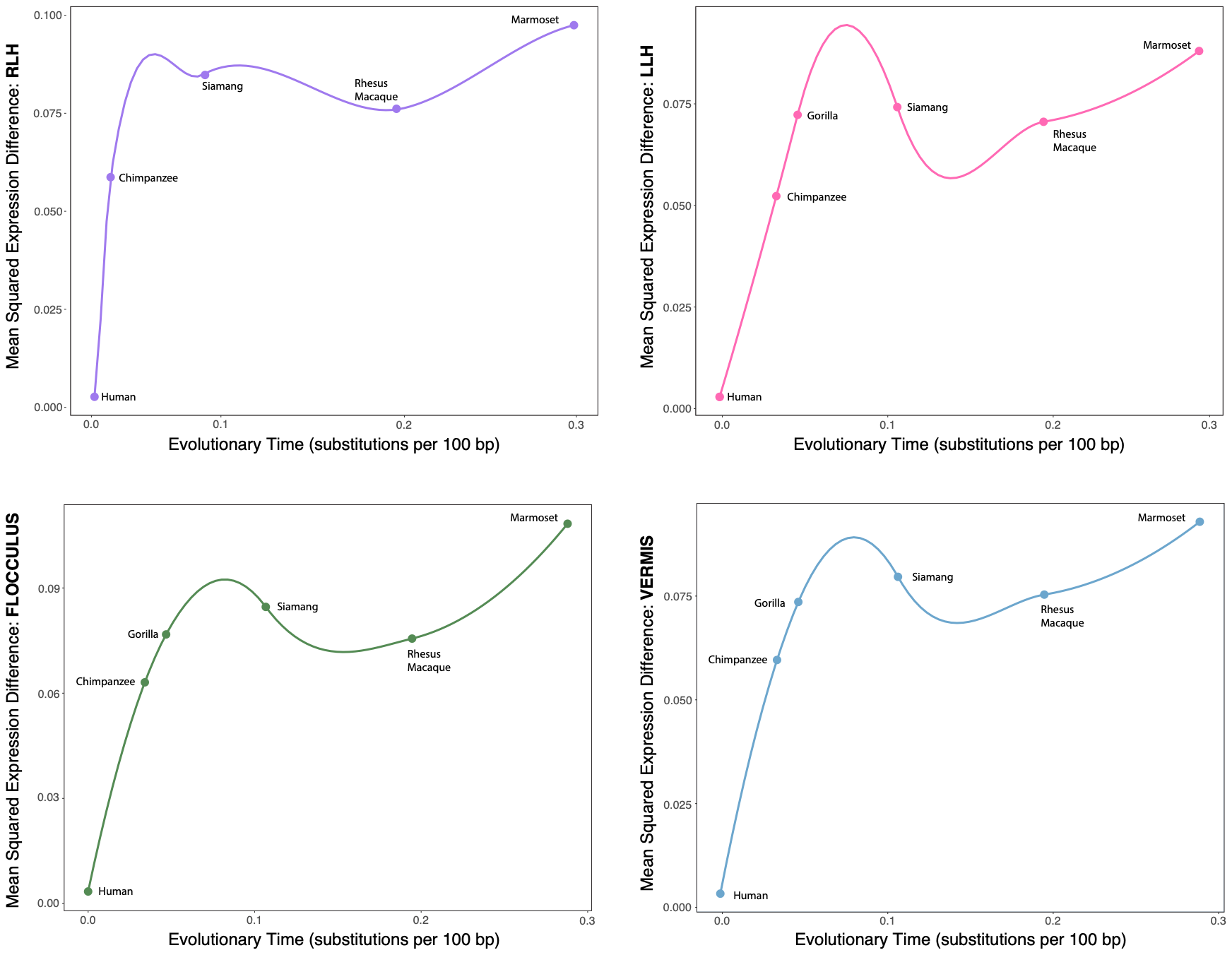

**Supplementary Figure 2:  Mean-square expression differences over evolutionary time for each of the cerebellar regions.** Each dot represents one of the 5 species. Evolutionary time is estimated by the number of substitutions that occur over a 100bp region. Mean squared expression difference was obtained using the residuals.R script from the EVEE-tools suite. The overall trend is summarized by the smooth line generated using a generalized linear model (GLM).

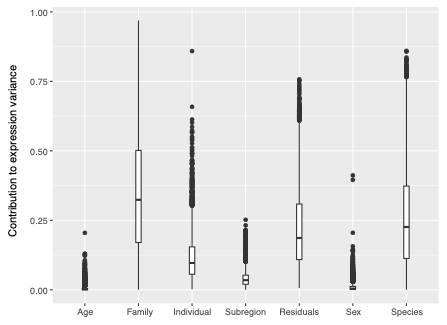

**Supplementary Figure 3: Boxplots of the distributions of variance contributions as determined through ANCOVA analysis.** The analysis evaluated the relative impact of age, family, individual variation, subregion, sex, and species, as well as residual variance representing unexplained variation. The y-axis indicates the proportion of total expression variance attributed to each factor. For age, we used the raw values and did not correct for lifespan. These relative contributions are for the total dataset, not separated by individual subregions.

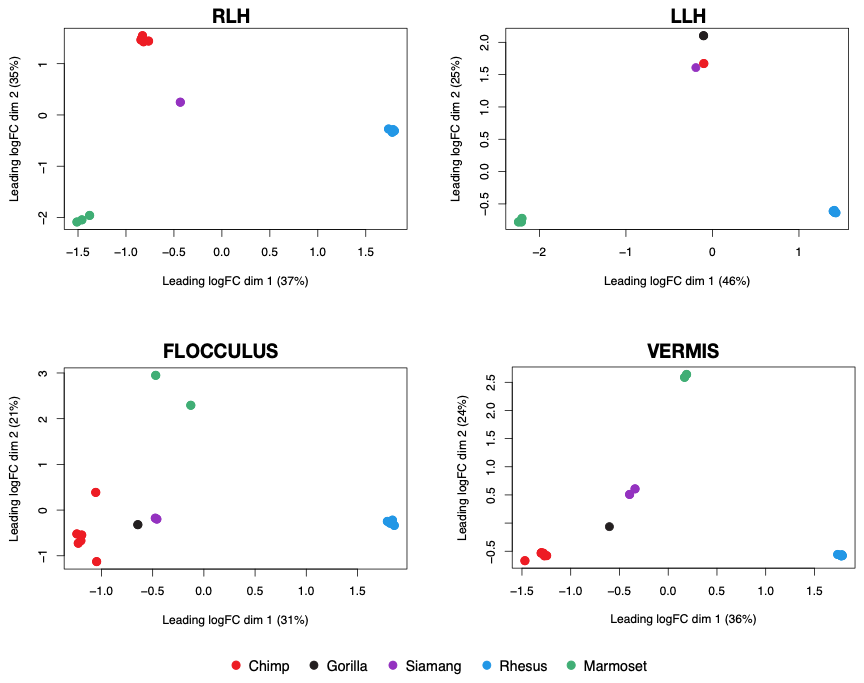

**Supplementary Figure 4: MDS plots by cerebellar subregion of non-human primate samples.** MDS plots were generated in EdgeR, using the plotMDS.DGElist function. Only the first two dimensions are shown. For non-human comparisons, the human samples were removed from the datasets and variation was recalculated based on only the remaining 5 (4 for the RLH) species. Each plot was calculated using the top 500 differentially expressed genes for each subregion. Samples in each plot are color-coded by taxa identity.

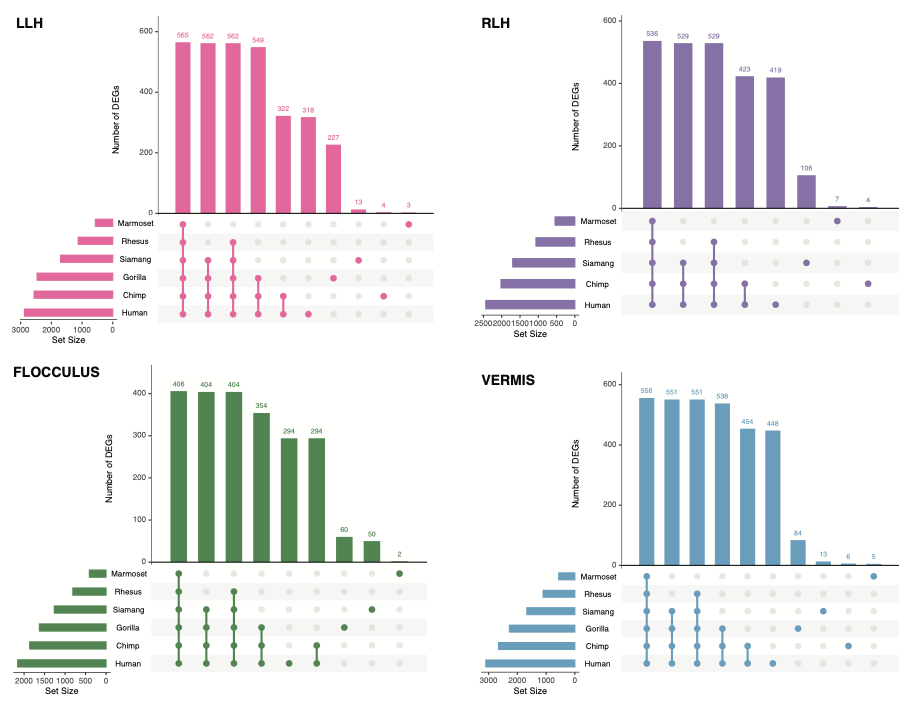

**Supplementary Figure 5: UpSet Plots of each subregion showing the shared DEGs for all species across the phylogeny.** DEGs were calculated using EVEE-tools and each species was treated as a single regime in order to assess which genes are uniquely expressed in particular species. The number of DEGs in each species grouping is depicted in the bar plot. The set size refers to the total number of genes associated with a given species.

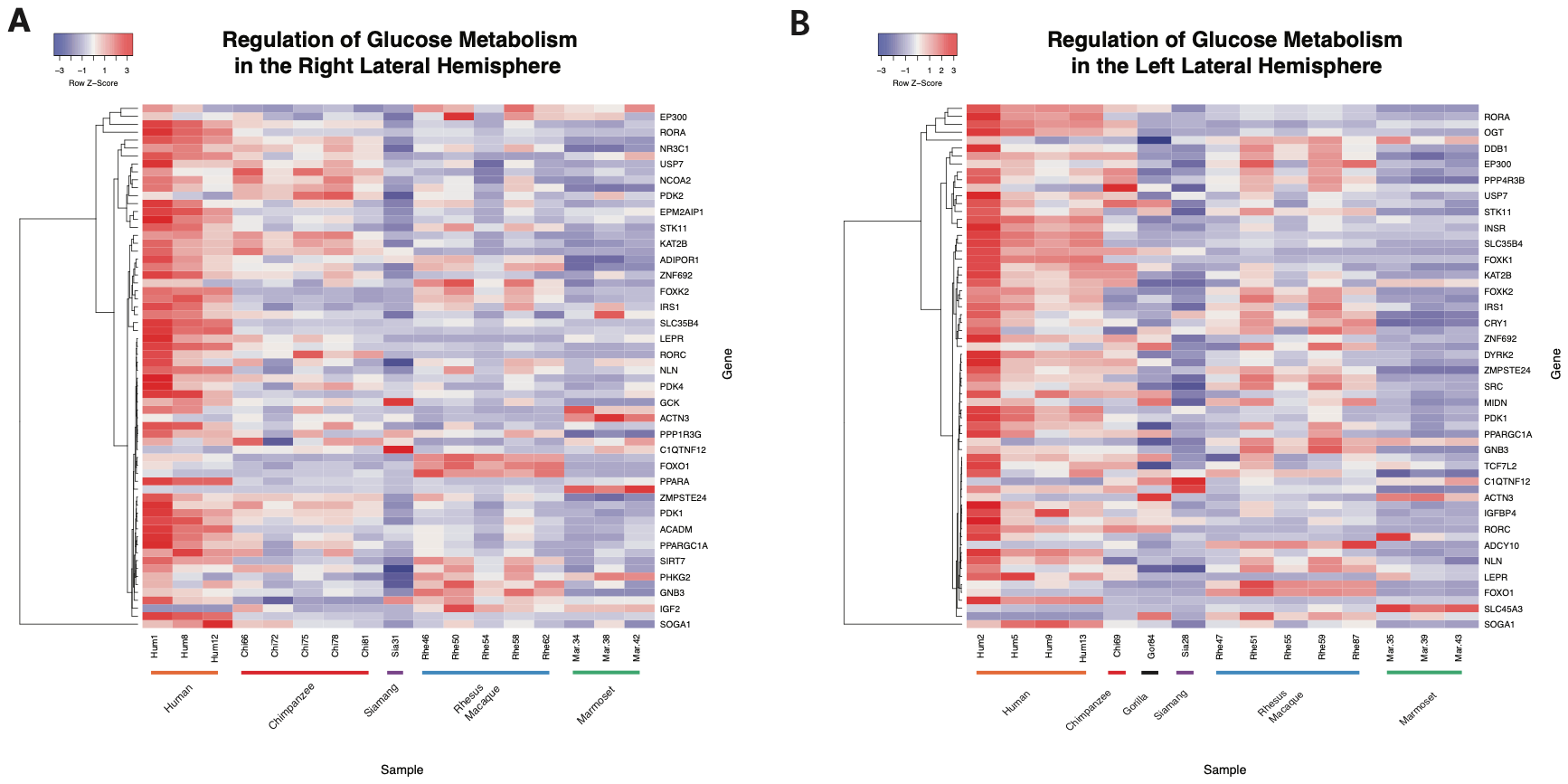

**Supplementary Figure 6: Heatmaps of individual lateral hemispheres of genes involved in the regulation of glucose metabolism (GO:0010906).** Each row represents a single gene from the GO set of terms. Each column represents a sample: samples are grouped according to species (labeled along the x-axis). Heatmap colors represent relative expression levels (red: more highly expressed; blue: lowly expressed). Heatmaps were generated using heatmapper.ca (Babicka et al. 2016).

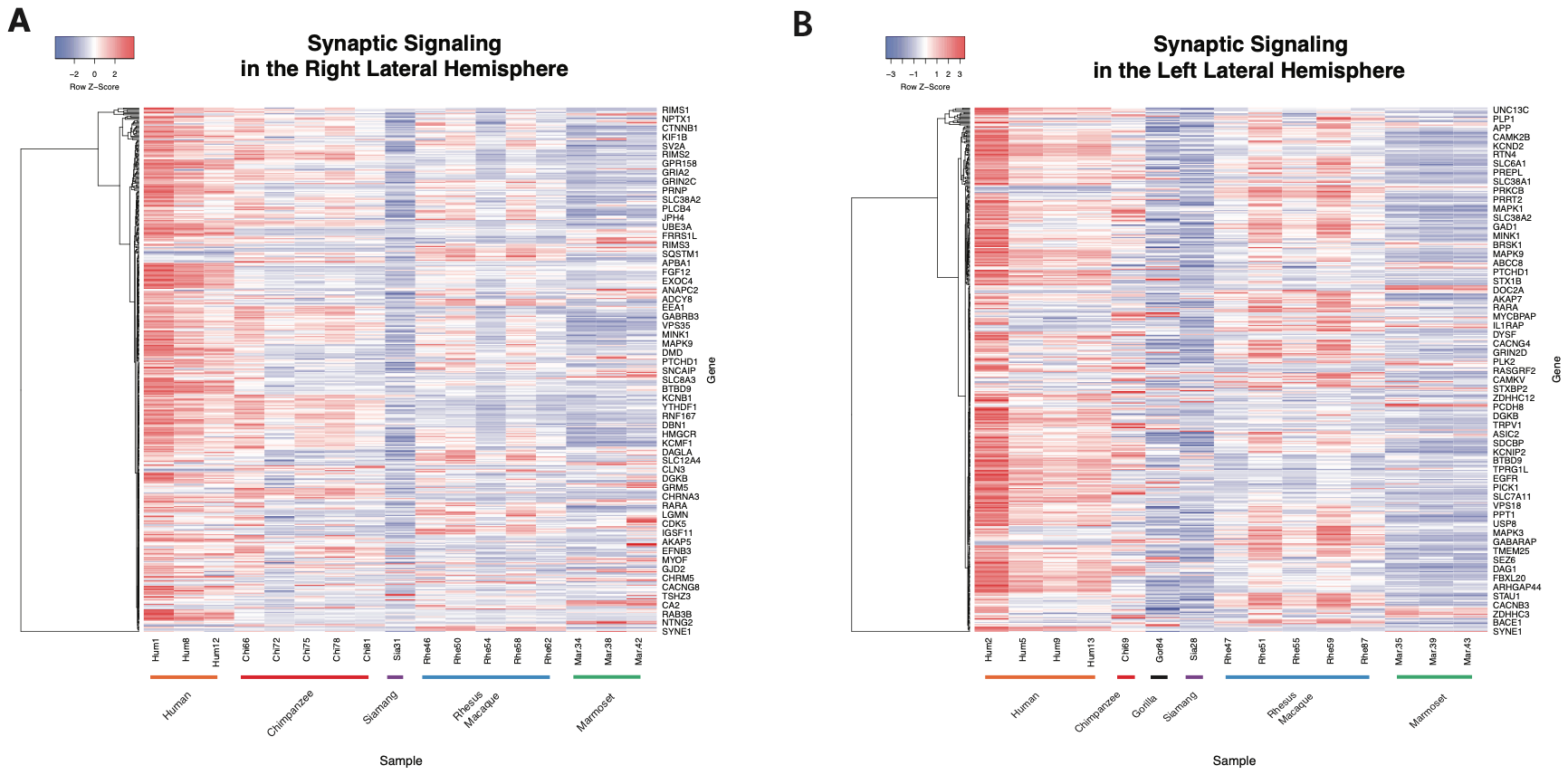

**Supplementary Figure 7: Heatmaps of individual lateral hemispheres of genes involved in synaptic signaling (GO:0099536).** Each row represents a single gene from the GO set of terms. Each column represents a sample: samples are grouped according to species (labeled along the x-axis). Heatmap colors represent relative expression levels (red: more highly expressed; blue: lowly expressed). Heatmaps were generated using heatmapper.ca (Babicka et al. 2016).

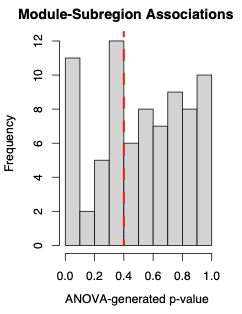

**Supplementary Figure 8: Histogram of p-values from ANOVA test for gene module association with cerebellar subregions.** We used the WGCNA package in R to find modules of co-expressed genes in each species. In order to assign these modules to particular cerebellar subregions, ANOVA tests were performed. The distribution of p-values from these ANOVA tests are depicted in a histogram. A significant cutoff at 0.4 is depicted by a red dotted line: the p-values to the left of the line represent a relatively normal distribution. Only module-subregion pairings with a p-value of 0.4 or less were considered based off of this distribution.

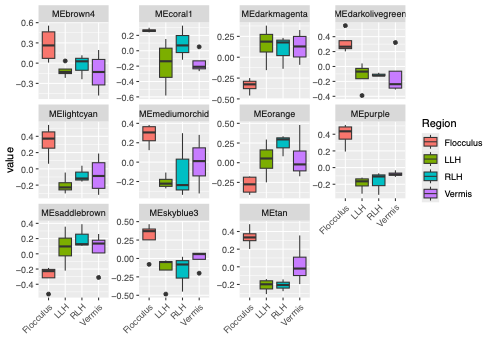

**Supplementary Figure 9: Boxplots of different human gene co-expression modules and their relative associations with each cerebellar subregion.** Each module of co-expressed genes was created using the WGCNA package in R. We performed ANOVA tests to pair each module with a cerebellar subregion. The y-axis represents the relative association between gene variation in a module and gene variation in a cerebellar subregion. Each plot represents a different module, and each box within the plot represents a different subregion (color-coded).
